# Microenvironmental engineering improves the self-organization of patterned microvascular networks

**DOI:** 10.1101/2022.04.04.487052

**Authors:** Katelyn A. Cabral, Vasudha Srivastava, Maxwell C. Coyle, Connor Stashko, Valerie Weaver, Zev J. Gartner

**Affiliations:** Graduate Program in Bioengineering, University of California, San Francisco and University of California, Berkeley; Department of Pharmaceutical Chemistry, University of California, San Francisco; Department of Molecular and Cellular Biology, University of California, Berkeley; Center for Bioengineering and Tissue Regeneration, Department of Surgery, University of California, San Francisco; Chan Zuckerberg Biohub, University of California, San Francisco; Center for Cellular Construction, University of California, San Francisco

## Abstract

The construction of three-dimensional (3D) microvascular networks with defined structures remains challenging. Emerging bioprinting strategies provide a means of patterning endothelial cells (ECs) into the geometry of 3D microvascular networks, but the microenvironmental cues necessary to promote their self-organization into cohesive and perfusable microvessels are unknown. To this end, we reconstituted microvessel formation *in vitro* by patterning thin lines of closely packed ECs fully embedded within a 3D extracellular matrix (ECM) and observed how different microenvironmental parameters influenced EC behaviors and their self-organization into microvessels. We found that the inclusion of fibrillar matrices, such as collagen I, into the ECM positively influenced cell condensation into extended geometries such as cords. We also identified the presence of a high molecular weight protein(s) in fetal bovine serum (FBS) that negatively influenced EC condensation. This component destabilized cord structure by promoting cell protrusions and destabilizing cell-cell adhesions. Endothelial cords cultured in the presence of fibrillar collagen and the absence of this protein activity were able to polarize, lumenize, incorporate mural cells, and support fluid flow. These optimized conditions allowed for the construction of branched and perfusable microvascular networks directly from patterned cells in as little as three days. These findings reveal important design principles for future microvascular engineering efforts based on bioprinting techniques.

## Introduction

Bioengineers can facilitate the ability of cells to self-organize into tissues by providing them with appropriate microenvironmental cues^1–3^. In the case of vasculature, many efforts have explored the microenvironmental cues that promote the formation of endothelial networks through a self-organizing process that mimics vasculogenesis^4–6^. Vasculogenesis typically begins with a randomly distributed population of cells and progresses through the coordination of a variety of distinct cell behaviors including proliferation, mechanical polarization, motility, adhesion stabilization, and lumenization^7^. However, vasculogenesis (and self-organization in general) is also a highly stochastic process^8^. Consequently, the patterns of microvessels formed are challenging to control. By employing techniques such as bioprinting and microscale engineering, bioengineers can provide additional spatial cues to guide self-organization more deterministically by reducing the subset of cell behaviors required to form mature microvessels, and thus, limiting the influence of stochastic processes on cell positioning^6,9^. Optimal deployment of these methods requires an understanding of the influence of each component of the microenvironment on the cell behaviors necessary for each stage of endothelial cell (EC) self-organization.

Strategies for engineering microvasculature can be broadly categorized into bottom-up and top-down approaches^9^. Bottom-up approaches aim to engineer the self-organization processes of vasculogenesis and angiogenesis. When ECs are seeded within an extracellular matrix (ECM), they self-organize into microvascular networks with a broad distribution of network geometries and vessel diameters^4,6,10,11^. With application of interstitial fluid flow, these capillaries can lumenize and become perfused^12–14^. However, this process takes up to 7 days to complete due to the slow speed of angiogenesis (5 µm/hour)^15^ and does not readily create hierarchical networks optimized for fluid transport^16,17^. Consequently, the center of thick and cell-dense tissues vascularized using these methods will suffer from hypoxia prior to completion of the process^18,19^. Top-down approaches, exemplified by bioprinting and microfabrication techniques, involve fabrication of hollow channels within hydrogels that are subsequently lined with a monolayer of ECs^20–25^. This strategy excels at creating vascular structures that are hundreds of microns to millimeters in diameter with predefined geometry. However, the creation of capillary-sized vessels is challenging with top-down approaches due to the resolution limits of bioprinting and the tendency of narrow channels to clog when perfused with cell suspensions^9,26^.

Some bioprinting techniques allow for the rapid and direct placement of cells within ECMs at a high density, representing a potential hybrid approach^3,27^. Placing cells into a structure that resembles a vascular network effectively reduces the number of cell behaviors necessary to complete the process of self-organization and mitigates the effects of stochasticity. This potentially leads to more rapid vasculogenesis and better recapitulates more native-like and hierarchical vessel network geometries that are optimized for fluid flow^28,29^. For example, Brassard et al. demonstrated the bioprinting of high-density cell solutions within Matrigel or collagen hydrogels^3^. Over time, the cells self-organized and condensed into a single cohesive structure in the geometry dictated by the initial print. However, a limitation of this and other bioprinting techniques are the requirements on nozzle geometry and cell density, which effectively limit the diameter of the extruded close-packed and continuous vascular cords to around 250 microns – considerably larger than the diameter of capillaries. Consequently, no studies have yet reported on the microenvironmental cues required for narrow caliber bioprinted vascular cords to self-organize into stable and perfusable networks.

To investigate the formation of microvessels less than 100 µm in diameter, we used DNA Programmed Assembly of Cells (DPAC), a high-resolution cell patterning technique that allows for the formation of cell-dense tissues, fully embedded in a biologically relevant ECM, and with near single-cellular resolution in X and Y. We created capillary-sized lines of ECs within a 3D ECM and observed their self-organization into microvessels under the influence of different microenvironmental cues. We found that by directly engineering the microenvironment, specific cell behaviors involved in latter stages of vasculogenesis can be encouraged, ultimately leading to the orderly self-organization of the cells into perfusable networks and across length scales that have not been previously achieved.

## Methods

### Cell culture

Human umbilical vein endothelial cells (HUVECs, Lonza) were cultured in EGM-2 and used between passages 4 and 6. Human brain vascular pericytes (HBVPs) were cultured in DMEM + 10% FBS + penicillin/streptomycin and used between passages 4 and 13. EGFP-HUVECs, EGFP-HBVPs, and mCherry-HUVECs were created by transducing cells with a pSicoR-EF1a-GFP or mCherry lentivirus. HUVECs with cytosolic EGFP and nuclear mScarlet were created by transducing with a pSicoR-EF1a-EGFP-H2B-mScarlet lentivirus. All lentiviruses were made by the UCSF Viracore. Transduced cells were sorted on a BD Aria II flow cytometer.

### Cell patterning

Cells were patterned using DPAC^30–32^. Briefly, aldehyde-functionalized glass slides (Nexterion) were micropatterned with amine-functionalized DNA oligonucleotides, either by spotting with a Nano eNabler (BioForce Nanosciences)^30,31^ or by photolithography^32,33^. Reductive amination with 1% sodium borohydride resulted in a stable amine linkage between the oligonucleotides and the slide. The slide was treated with (tridecafluoro-1,1,2,2-tetrahydrooctyl) dimethylchlorosilane (Gelest) to render it hydrophobic and reduce non-specific cell adhesion. PDMS flow cells were plasma oxidized (Plasma Etch), placed atop the patterned slides, and treated with 1% BSA to further block non-specific cell adhesion.

A cell suspension of HUVECs in serum-free EGM-2 was incubated with lipid- or cholesterol-modified oligonucleotide complexes complementary to the oligo patterned on the slide^30–32^. After washing to remove excess oligo, the cells were re-suspended at a concentration of 30 million cells/mL, added to the PDMS flow cells, and passed over the patterned slide. Washing out excess cells revealed selective adhesion of the cells to the DNA pattern. Multi-layered endothelial cords were created by repeating this process with cells bearing oligonucleotides complementary to those in the previous layer.

A hydrogel precursor solution containing 2% Turbo DNase (Thermo Fisher Scientific) was added to the inlet of the PDMS flow cell and allowed to set around the patterned cells for 45 minutes at 37°C. The cell-containing hydrogel was then carefully removed from the slide and placed atop a drop of liquid hydrogel precursor solution. This process fully embedded the 2D-patterned cells into a 3D microenvironment. After 45 minutes of incubation at 37°C, media was added and forceps were used to remove the PDMS flow cells from the hydrogel.

### ECM hydrogel

Growth factor reduced Matrigel (Corning) was mixed with 2% Turbo DNase and neutralized rat tail collagen I (Corning) for a total protein concentration of 8 mg/mL. The ratio of Matrigel to collagen varied by experiment. Non-specific staining of the ECM was achieved by mixing the hydrogel precursor solution with 1% Alexa Fluor 647 NHS Ester (Invitrogen).

### Media

Endothelial cords were cultured in EGM-2 (EBM-2 with the addition of EGM-2 BulletKit) (Lonza). For most experiments, 50 ng/mL phorbol-12-myristate-13-acetate (PMA) (Sigma-Aldrich) was added to the media. For serum-free media, the FBS was omitted from the EGM-2 (normally 2% of the volume). Recombinant human VEGF-A, FGF-basic, TGF-β1, stem cell factor (SCF), stromal cell-derived factor-1α (SDF-1α), and interleukin-3 (IL-3) were purchased from R&D Systems.

Multiple lots and formulations of FBS were evaluated, including FBS from EGM-2 Bulletkits (Lonza), UCSF Cell Culture Facility, and charcoal/dextran stripped FBS (Gemini Bio). Heat-inactivation of FBS was achieved by heating to 56°C for 30 minutes. The FBS was fractionated by centrifuging through molecular weight cutoff filters (Sterlitech). The resulting concentrate and filtrate were then diluted with PBS to the same volume as the FBS before filtration. To hydrolyze protein, 100 µg/mL of proteinase K (Fisher Scientific) was added to the FBS and then incubated at 55°C for 1 hour before being neutralized with 100 µg/mL PMSF (phenylmethylsulfonyl fluoride, Millipore Sigma). Exosomes were purified from FBS using a Total Exosome Isolation Reagent Kit (Invitrogen).

### Immunofluorescent staining

Tissues were fixed with 2% PFA for 45 minutes, permeabilized for 15 minutes with 1% Triton X-100 (Sigma-Aldrich) and blocked overnight with 10% goat serum. To avoid cross-reactivity with the Matrigel, tissues that would later be treated with anti-mouse antibodies were also blocked with AffiniPure Fab Fragment Goat Anti-Mouse IgG (Jackson Laboratories). A list of antibodies used can be found in the Supplementary Methods.

### Time-lapse microscopy

Patterned cell pairs were incubated for 12 hours before being placed onto a Zeiss LSM800 microscope and imaged with a 25x objective every hour for 48 hours. The cells were fed with media supplemented with 50 mM HEPES and ProLong Live Antifade Reagent (Thermo Fisher).

### Image quantification

Cord morphology was quantified using FIJI^34^. The continuous length of each segment of cord were traced by hand (visualized by cytoplasmic EGFP). Tip cells were identified based on their orientation (perpendicular to the patterned cord) and the presence of long, thin subcellular protrusions. Disconnected cells that had migrated away from the cord were also included in the tip cell count as we had observed that the cells branching out from the cord tended to detach and migrate over time. The number of protrusions per cell at 60 hours of culture was counted manually. Cell protrusions were defined as extensions of the cytoplasm that were narrower than the nucleus and at least two microns in length. The number of frames that the cells of each pair were in contact was manually counted.

### Statistics

All statistical analyses were performed using Prism 9. For analyses comparing two experimental conditions, an unpaired t-test was performed. For analyses comparing three or more experimental conditions, a one-way analysis of variance (ANOVA) was used to compare the means, followed by Dunnett’s multiple comparisons test.

### Perfusion

Aluminosilicate glass micropipettes with a long tether were prepared using a P-97 micropipette puller (Sutter Instruments). The pulled pipettes were cut 3-5 mm from the tip to get 10-25 µm diameter pipettes with jagged ends. The pipettes were filled with PBS or PBS containing 2 µm-diameter Fluospheres Carboxylate-Modified Microspheres (Thermo Fisher). The pipette was mounted on a Narishige MM-89 micromanipulator connected to a syringe. The endothelial cords were cut on one end to create an opening and the micropipette was used to puncture the opposite end of the cord. Once the tip of the micropipette was within the lumen of the cord, the syringe created a positive pressure and induced flow of liquid and debris towards the open end of the cord. Images were acquired using an Axiovert 200M epifluorescence microscope.

## Results

We developed a reproducible assay to study the effect of microenvironmental cues on prepatterned endothelial cells by constructing cell-dense, 35 µm-wide, 2 millimeter-long lines of HUVECs using DPAC (Figure 1A)^30,32^. DPAC allows for the placement of cells with single-cell precision in a common imaging plane and fully embedded within a 3D hydrogel. Using DPAC, we reproducibly patterned capillary-sized lines of ECs with an average nucleus-to-nucleus spacing of 25 µm, compared to ∼15-21 µm nuclear spacing of capillaries *in vivo*^35^ (Figure 1B).

**Figure 1:**
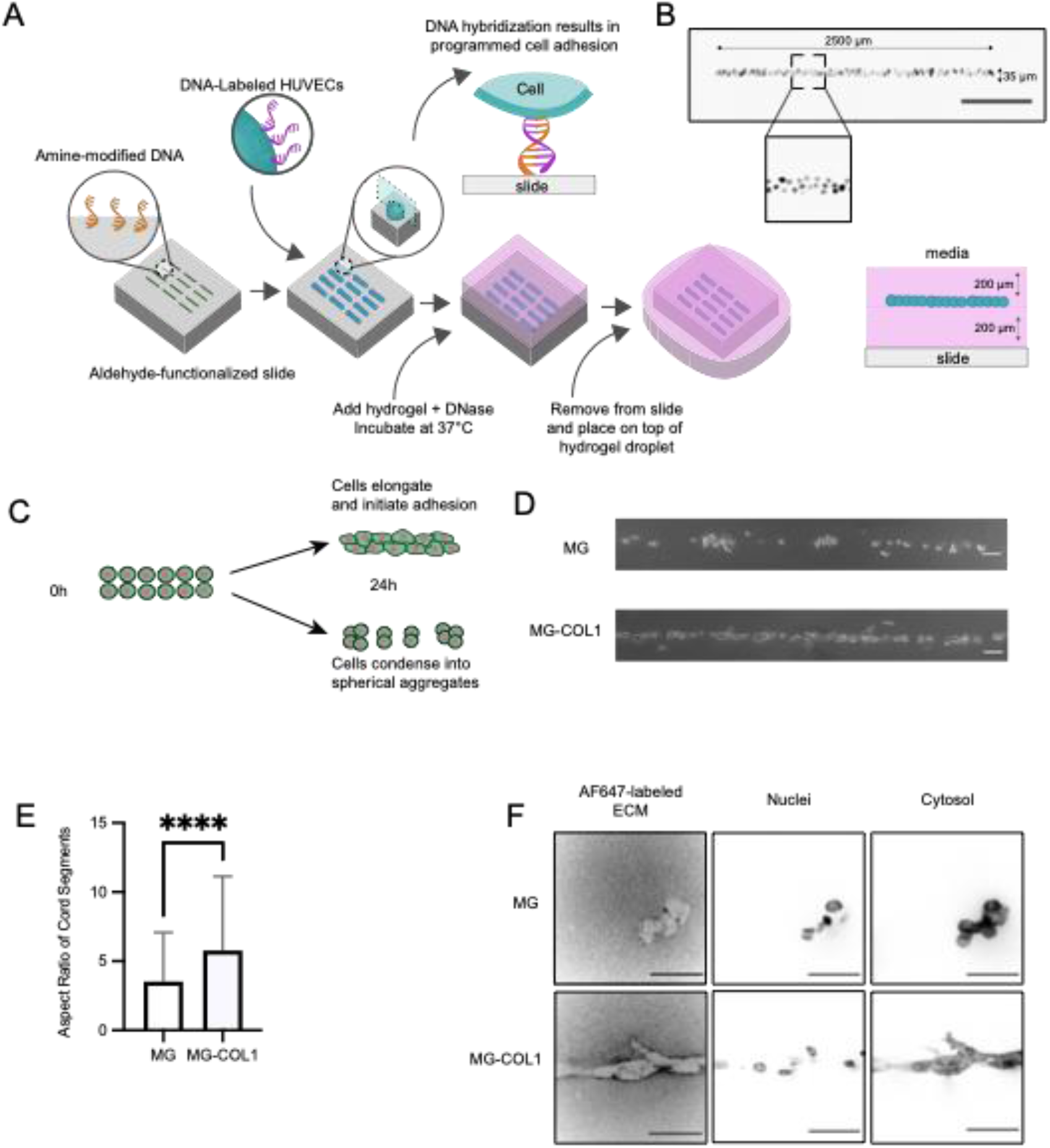
DNA-Programmed Assembly of Cells (DPAC) creates capillary-sized lines of HUVECs to assess the effect of ECM composition on EC self-organization. A) Diagram of the cell patterning process. First, photolithography is used to selectively expose regions of an aldehyde-functionalized slide^33^. An amine-modified DNA oligonucleotide solution is then dropped onto the surface. After baking and reductive amination, the amine-modified DNA is covalently bonded to the slide in the pre-defined pattern. HUVECs are then labeled with a set of lipid- or cholesterol-modified complementary oligos that insert into the cell membrane. Upon passing the cells over the surface, the DNA hybridizes, adhering the HUVECs to the surface only in the regions defined by the DNA pattern. The HUVECs are then embedded in a hydrogel containing 2% DNase. After gelation, the hydrogel containing the patterned HUVECs is removed from the slide and placed on top of another droplet of hydrogel. The result is HUVECs that have been patterned with high precision in a single 2D plane but embedded within a 3D ECM. B) Images of patterned HUVEC cords immediately after being fully embedded within the hydrogel. Scale bar: 500 µm. C) Diagram showing the condensation of patterned HUVECs into either a single continuous structure or several smaller structures. D) After 24 hours of culture, patterned lines of HUVECs in MG broke up into several smaller, rounded sections (top). Patterned lines of HUVECs in MG-COL1 hydrogels were able to condense into a single cord structure (bottom). Scale bar: 100 µm. E) The aspect ratio of cord segments cultured in MG-COL1 were significantly increased (T-test, p<0.0001). F) HUVEC cords with fluorescent nuclei (H2B-mScarlet) and cytoplasm (EGFP) were cultured in either MG or MG-COL1 that had been stained with an Alexa Fluor 647 NHS-ester dye to visualize the ECM.

We first investigated components of the ECM microenvironment that promote the coalescence of micropatterned HUVECs into long multicellular cords (Figure 1C), focusing on combinations of basement membrane proteins (e.g., Matrigel) and stromal ECM (e.g., collagen I). We found that EC lines cultured for 24h in growth factor reduced Matrigel (MG) were mechanically unstable, fragmenting and then condensing into several cell “droplets” (Figure 1D). In contrast, identical EC lines cultured in a composite ECM of 6 mg/mL MG and 2 mg/mL collagen I (MG-COL1) were better able to maintain an extended morphology over 24h (Figure 1E). Controlling for total protein concentration (8 mg/mL), we found that increasing the COL1 concentration in the hydrogel increased the length of the cord segments (Figure S1A). We considered the possibility that COL1-containing gels increased endothelial spreading and angiogenic outgrowth due to elevated matrix stiffness^36,37^. However, there was no significant difference in the elastic modulus (E) between 8 mg/mL MG and MG-COL1 hydrogels, as measured by bulk hydrogel rheology (Figure S1B) and atomic force microscopy (Figure S1C). An alternative explanation stems from the ability of cells to mechanically remodel their microenvironment, including synthesizing ECM proteins, concentrating ECM components, and aligning them anisotropically^38–40^. We examined matrix remodeling using fluorescently labeled matrices. EC lines cultured in MG-COL1 gels concentrated ECM into bright sheaths at the cord surface, perhaps providing a more mechanically stable substrate to maintain an extended morphology (Figure 1F)^39^. The same was not true for EC lines cultured in Matrigel.

While collagen-containing hydrogels support initial EC elongation into cord-like structures over 24 hours (Figure 2A), they failed to maintain this architecture and mature after extended culture. Instead, we found that the cells scattered over the following 48 hours (Figure 2B). During this process, many cells exhibited tip cell-like morphology with numerous subcellular protrusions^41,42^. We hypothesized that EGM-2 media lacked factors necessary to maintain and stabilize cell-cell junctions. PMA has been previously shown to promote angiogenesis^43–45^. Addition of 50 ng/mL PMA modestly reduced HUVEC scattering and increased the length of cord segments (Figure S2A). The addition of pro-angiogenic, pro-vasculogenic growth factors^46–48^ FGF-2, VEGF-A, and TGF-β had no effect on cord morphology (Figure S2B). The combination of SDF-1*α*, IL-3, and SCF has been shown previously to promote vascular tube morphogenesis^47^ but had no effect in this assay (Figure S2B).

**Figure 2:**
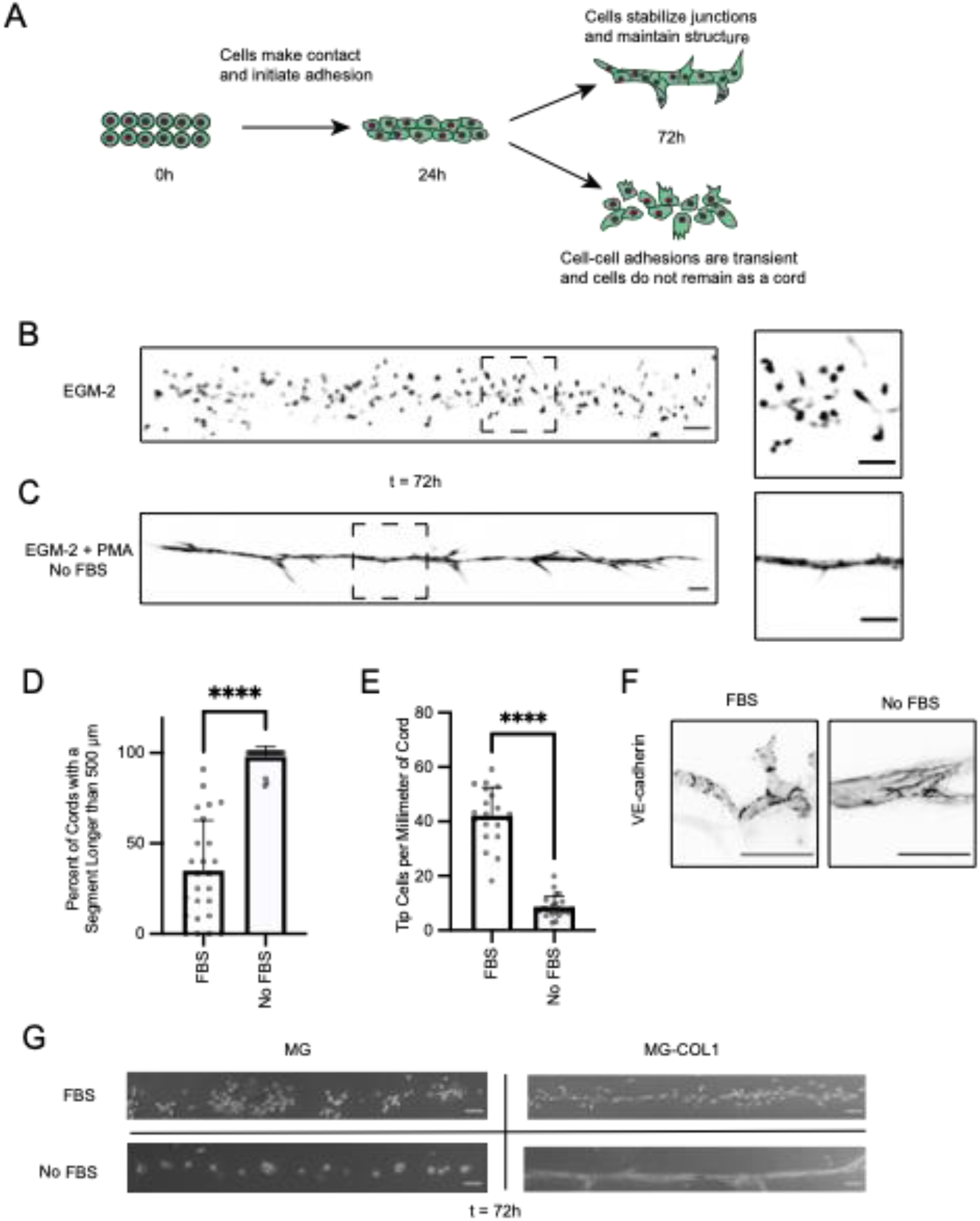
Addition of PMA and removal of FBS results in stable, cohesive HUVEC cords with continuous adherens junctions. A) Diagram showing that after the initial condensation of the patterned HUVECs into a single continuous structure, we need to find microenvironmental conditions to maintain that structure and stabilize the cord. B) After 72 hours of culture in EGM-2 media, HUVEC cords embedded in MG-COL1 scattered into small segments and individual cells. C) After 72 hours of culture in EGM-2 media without FBS, the HUVEC cords maintained one cohesive structure in the approximate shape of the original patterned vessel. D) The percent of cords that had at least one segment longer than 500 µm in length at 72 hours of culture in EGM-2 + PMA that either contained or omitted FBS. Bars represent mean ± SEM and with each data point representing one independent tissue. T-test, p<0.0001. E) The number of angiogenic sprouts per millimeter of HUVEC cords cultured for 72 hours in either FBS-free EGM-2 + PMA or FBS-containing EGM-2 + PMA. Data represented as mean ± SEM and with each data point representing one independent tissue. T-test, p<0.0001. F) VE-cadherin staining in HUVEC cords cultured in either FBS-free EGM-2 + PMA or FBS-containing EGM-2 + PMA. Scale bar = 50 µm. G) Comparison of HUVEC cords grown in either MG or MG-COL1, and in media containing or omitting FBS, after 72h of culture.

The inability to stabilize cord morphology through added factors suggested an alternative hypothesis — that EGM-2 contained factors that antagonize cell behaviors necessary for stable cord formation. Fetal bovine serum (FBS) is a component of EGM-2 that contains many poorly characterized factors that could contribute to cell scattering^49,50^. Strikingly, omitting FBS, normally 2% of the volume of EGM-2, had a dramatic effect on cord morphology. Instead of scattering, the HUVECs remained tightly adhered to one another and maintained the original patterned shape (Figure 2C, 2D). There was also a reduction in the number of tip cells arising from the main cord (Figure 2E). Staining for VE-cadherin revealed that adherens junctions are impacted by FBS (Figure 2F), having a ragged appearance at cell-cell junctions and exhibiting punctate staining within the cell, consistent with endocytosis and an angiogenic phenotype^51,52^. These results suggest that FBS encourages a pro-angiogenic phenotype, leading to the destabilization of adherens junctions and breakdown of endothelial cord structure. Combining these findings with the stabilizing influence of COL1-containing ECM, we concluded that matrices containing fibrillar collagen promote more extended morphologies than pure Matrigel and that factors within FBS destabilize a cord-like morphology (Figure 2G).

FBS contains many beneficial components that support cell viability and growth^49^. We therefore sought to identify whether we could selectively remove the activity in FBS contributing to the scattering phenotype without omitting the beneficial components. Fractionation of the FBS using molecular weight cutoff filters demonstrated that the active component(s) of FBS had a molecular weight greater than 200 kDa (Figure S3). The activity was present in heat-inactivated FBS, charcoal-stripped FBS, and exosome-depleted FBS, suggesting it was not part of the complement system, a lipophilic molecule, or an exosome, respectively. Proteinase K treatment of the FBS blocked its effects, suggesting that the active component of FBS contains a protein. We tested a range of candidate proteins that are known to be present in plasma or serum^50,53^ (Figure S4) but none were sufficient to induce cell scattering from patterned cords when added to the media.

To better understand how the unidentified component in FBS was contributing to cord destabilization, we used live cell imaging to examine individual cellular behaviors within the patterned cord. In the presence of FBS, we observed ECs extending many protrusions, making transient adhesions with other ECs, and then migrating away from the main cord (Video S1). In the absence of FBS, the ECs were elongated and moved back and forth along the cord but remained adhered to other ECs (Video S2). To investigate these cell-cell interactions in greater detail, we used DPAC to pattern pairs of ECs separated by 25 µm as minimal “tissues” that were cultured in the presence and absence of FBS (Figure 3A). Time-lapse microscopy revealed striking differences in morphology and cell behavior due to the media composition (Videos S3, S4). EC pairs cultured with FBS had a star-shaped morphology, with protrusions extending in all directions. In contrast, EC pairs cultured in serum-free media tended to elongate and had fewer protrusions (Figure 3B). FBS also resulted in more transient cell-cell adhesions, as measured by the fraction of time where the cells were in contact as well as the average number of instances where the cells separated over the course of the time-lapse (Figure 3C). These observations further suggest that the active component in FBS destabilizes cords through its effects on cell-cell junctions and by promoting cell morphologies and patterns of motility that increase the probability of cells moving away from their nearest neighbors.

**Figure 3:**
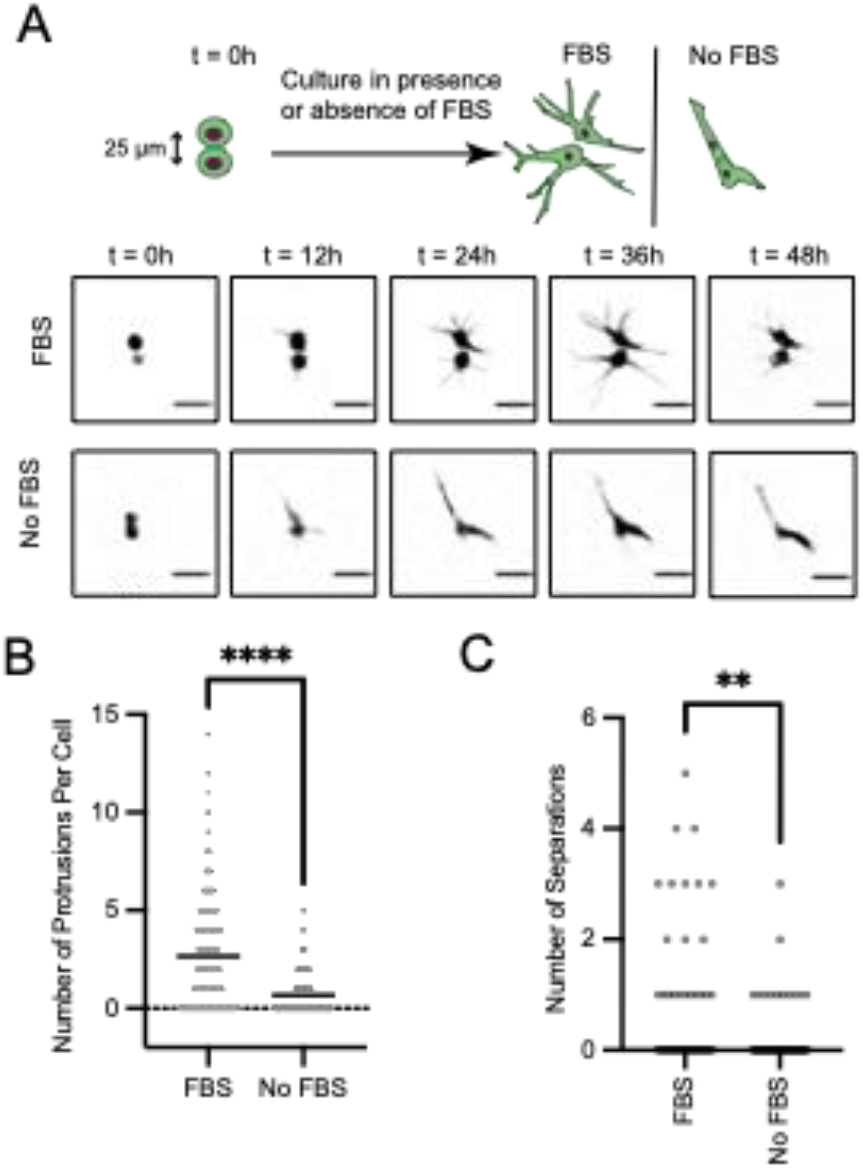
FBS increases proliferation and extension of subcellular protrusions but inhibits the formation of stable cell-cell adhesions. A) Pairs of HUVECs were patterned 25 µm apart (center to center distance) and cultured for 60 hours in either FBS-free EGM-2 + PMA or FBS-containing EGM-2 + PMA. Cell pairs were imaged once per hour between hours 13-60. B) Quantification of the number of protrusions per cell. Bars represent the mean ± SEM. Each data point represents one cell. T-test, p<0.0001. C) Quantification of the average number of instances over the time-lapse where the two cells in a pair lost contact with each other. Bars represent the mean ± SEM. Each data point represents one cell pair. T-test, p<0.01.

Having identified microenvironmental conditions and cell behaviors that promote stable cord formation, we investigated how these structures progress through additional steps of microvascular self-organization (Figure 4A). Endothelial cords cultured without FBS were stable for at least seven days, exhibiting little change in cord morphology (Figure 4B). Staining for ZO-1 and claudin-5 revealed tight junctions between the ECs of these microvessels (Figure 4C, 4D). The presence of an endothelial-derived basement membrane around the endothelial cords was confirmed by staining for human collagen IV (Figure 4E).

**Figure 4:**
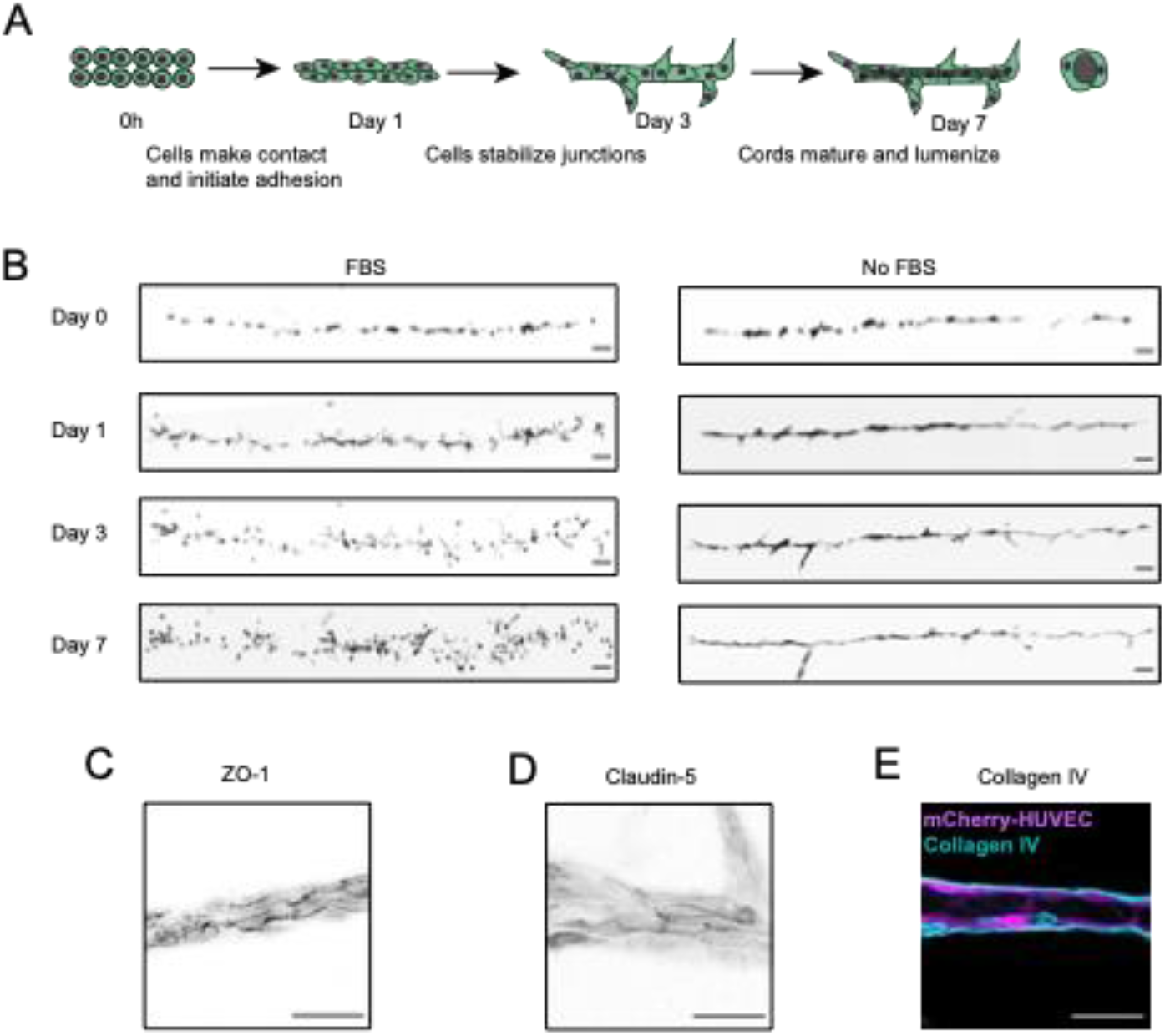
Formation of stable, lumenized, and mature microvascular networks of controlled dimensions after extended culture of HUVEC cords in serum-free media. A) Diagram of the HUVEC cords progressing through the stages of vasculogenesis towards maturation and lumenization. B) HUVEC cords were cultured in EGM-2 + PMA either containing or omitting FBS for 7 days. The same cords were imaged each time, demonstrating the extent of cord morphology change over time. Scale bar: 100 μm. C) ZO-1 localized to the cell-cell junctions. Scale bar: 50 μm. D) The endothelial tight junction protein claudin-5 was present at cell-cell junctions. Scale bar: 50 μm. E) HUVEC cords (magenta) were able to synthesize a basement membrane around themselves (collagen IV, cyan). Scale bar: 50 μm.

While cords appeared stable in this assay, we did not see clear evidence of lumenization - a final step in cord self-organization necessary for perfusion. Lumenization of blood vessels can occur through several different mechanisms^54–56^. We hypothesized that by increasing the cell-cell surface area, we could facilitate the formation of a lumen between ECs^57^. We patterned three-layered HUVEC cords that were 30-75 µm in diameter (Figures S5A, S5B) and condensed into a stable, continuous structure in serum-free media (Figures S5C). However, we also observed cell debris accumulating in the center of these larger cords consistent with the onset of lumenization by cavitation^58^ (Figure 5A). We investigated whether these vessels were perfusable by cutting the vessel at one end and piercing the other end with a micropipette connected to a syringe. Upon injection of PBS, the cell debris was flushed through the cord, revealing a fully lumenized, perfusable microvessel (Figure 5B, Video S5). Fluorescent microbeads injected into the lumen traveled the length of the vessel (Figure 5C, Video S6). The microvessels could be perfused 3-7 days after patterning. Confocal imaging revealed an open lumen and a polarized endothelium with apical actin and basal laminin α5 (Figure 5D). When human brain vascular pericytes (HBVPs) were patterned on top of the ECs, they formed close associations with the ECs and were embedded within the vascular basement membrane (Figure 5E). We created a variety of branched microvascular network structures (Figure 5F, Figure S6) which could also be perfused across branch points (Figure 5G, Video S7). Thus, by engineering the microenvironment to control the process of vascular self-organization from patterned ECs, we were able to create mature and perfusable microvessels incorporating mural cells with a controlled branching architecture.

**Figure 5:**
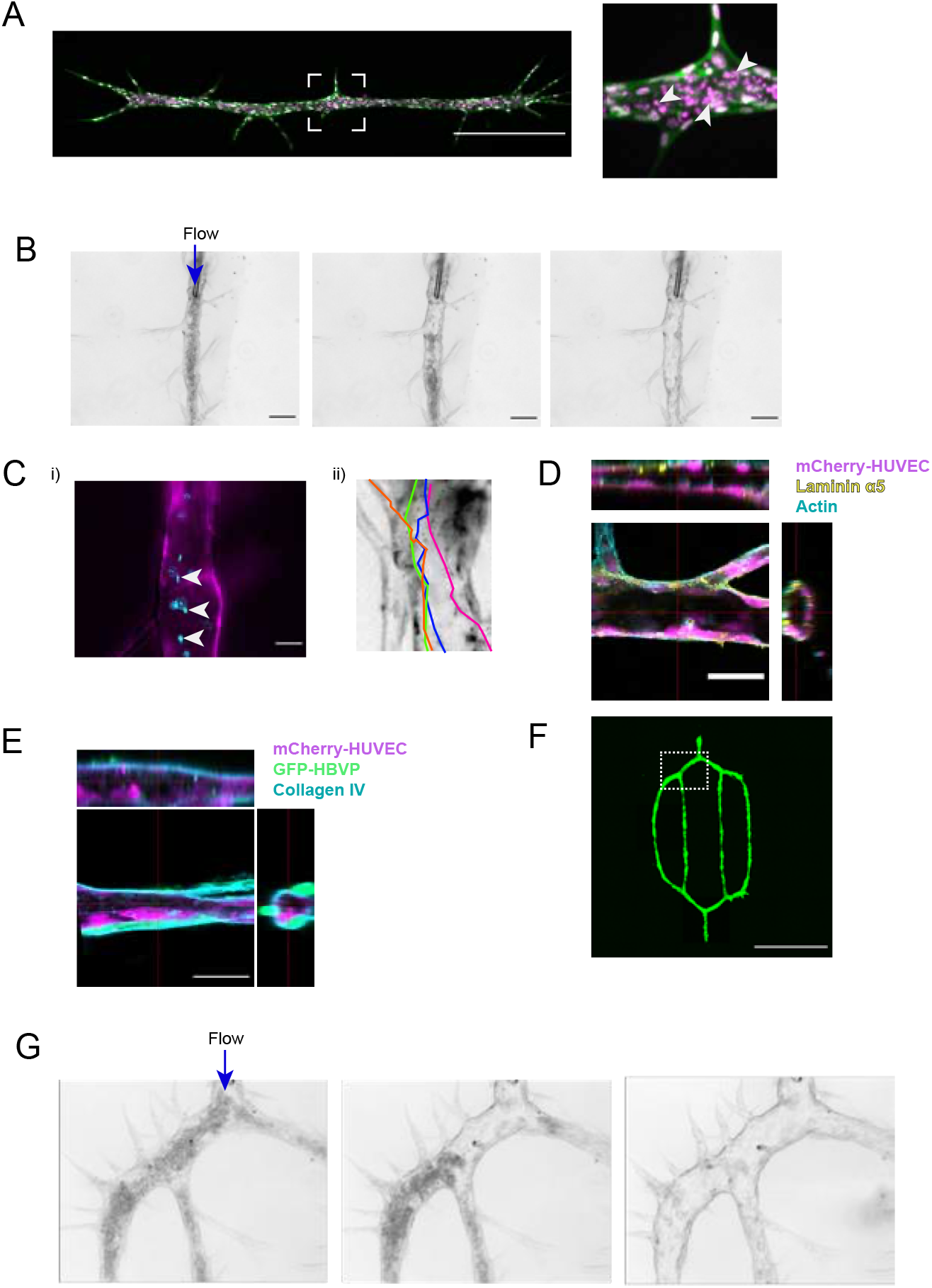
Microenvironmental engineering of lumenized and perfusable microvasculature with defined vessel architecture. A) Cell and nuclear debris (white arrowheads) observed in the center of a three-layered EGFP-EF1a (green), mScarlet-H2B (magenta) HUVEC cord after 5 days of culture in serum-free EGM-2. Scale bar: 500 µm. Inset: Closeup image of debris. Scale bar: 50 µm. B) Microinjection of PBS into a three-layered microvessel resulted in the removal of cell debris, revealing an intact lumen down the length of the microvessel. C) A three-layered mCherry-HUVEC microvessel (magenta) was perfused with 2 µm diameter blue fluorescent beads (cyan) (white arrowheads) Scale bar: 50 µm. D) A perfused cord was imaged with a confocal microscope, revealing an empty lumen surrounded by mCherry-HUVECs (magenta). Polarization is evident by actin (cyan) and laminin α5 (yellow) staining. E) A three-layered mCherry-HUVEC cord (magenta) was co-cultured with EGFP-HBVPs (green), leading to the deposition of a collagen IV (cyan)-rich basement membrane. F) A stylized branched microvascular network was created by patterning EGFP-HUVECs and culturing them in MG-COL1. Image taken after 24 hours of culture. Boxed region shown in detail in panel G. G) After 72 hours of culture, the branched structure was perfused with a micropipette. The images show a progression of the debris being washed out of the lumen.

## Discussion

Numerous studies have demonstrated efficient formation of microvessels through vasculogenesis, but control over the resulting network structures and vessel diameters is often lacking. We reasoned that tissue engineers could promote the maturation of prepatterned ECs into cords and vessels that retain fidelity to their designed pattern by considering the “design principles” of self-organization during vasculogenesis. Self-organization during traditional in vitro vasculogenesis assays involves a variety of EC behaviors that play specific roles during each stage of the process. ECs must first migrate, proliferate, and extend protrusions to establish contacts with other ECs, then switch to a different set of cell behaviors that promote maturation by stabilizing nascent cell-cell adhesions, polarizing, and lumenizing. By using high-resolution cell patterning to place the ECs adjacent to each other, the process effectively starts at the midway point, reducing the cell behaviors necessary for efficient self-organization. We used DPAC to test this idea by patterning ECs in cords of similar diameter and cell density to capillaries. These cords provided us with a reliable assay to test the effects of microenvironmental cues necessary for self-organization of patterned ECs into mature into perfusable microvessels.

Endothelial cord formation requires that initially close-packed cells stabilize cell-cell interfaces while maintaining an extended geometry on their basal interface. This process requires reorganization of the contractile apparatus at cell-cell and cell-ECM interfaces, and therefore requires a substrate capable of sustaining the resulting active tensions generated by the cells. We found that matrix composition is critical for this process. Patterned HUVEC cords cultured in Matrigel were unable to maintain an extended and cohesive structure. Instead, the patterned lines of cells broke up into several smaller cohesive clusters, or “droplets,” consistent with a high interfacial tension between the basal surface of the ECs and the surrounding gel^59,60^ (Figure 2G). We hypothesized that to balance their interfacial tension at the basal surface, the cells required a fibrous component of the ECM which can be organized into a more mechanically stable substrate. This concept plays an important role in the morphogenesis of a variety of tissues^39^. Indeed, we found that fluorescently labeled matrix was concentrated at the basal surface of HUVEC cords cultured in MG-COL1 through a process known to involve cell contractility^15,39^.

After the formation of cell-cell and cell-ECM interfaces, the microenvironment must provide signals supporting the maintenance of those interfaces while removing signals that promote interface turnover^61^. However, we found that groups of ECs cultured in Matrigel and collagen I disbanded into individual ECs after 72 hours of culture. We explored the addition of media factors previously described to promote cord formation through randomly seeded vasculogenesis or angiogenesis (e.g., VEGF, FGF)^47^. Surprisingly, these conditions did not enhance cord stability when the cells were seeded directly into a cord-like geometry. A potential explanation for this observation is that pro-angiogenic factors are useful when tip cell-like behaviors such as migration and extension of protrusions are desired^41,62^. However, when cell behaviors promoting microvessel stabilization and maturation are required, these factors provided no additional benefit and may in fact oppose microvessel stability. We reasoned a subset of factors present in FBS were promoting tip cell-associated behaviors, leading to destabilization of cell-cell interfaces. Careful examination of the underlying cell behaviors promoted by FBS revealed the extension of protrusions and a decreased lifetime of cell-cell interfaces, both behaviors associated with angiogenic tip cells but not stable microvasculature^63,64^. Removal of a high molecular weight and protein-based component of FBS allowed the engineering of stable, branched, and perfusable vascular networks. Thus, both the mechanical and chemical components of the microenvironment must be tailored to promote the specific cell behaviors necessary for the self-organization of prepatterned tissues—behaviors that might be opposed by factors necessary for cell expansion and migration.

The flexibility, scalability, and high-resolution of DPAC allowed us to create microvessels of diverse shapes and sizes, including capillary-sized microvessels (diameter ∼10 µm) and branched vascular networks millimeters in length. However, the microenvironmental engineering principles identified here should be applicable to other techniques capable of depositing ECs into controlled geometries fully embedded in ECM. Additionally, opportunities exist to combine this approach with other advanced methods for engineering microvasculature. For example, the microenvironmental cues identified in this paper could be applied to further promote the rapid self-organization of ECs patterned via extrusion bioprinting^3^. The strategies explored here could also be used to engineer narrower gauge vessels that interface with larger vessels prepared using biofabrication methods such as laser ablation^25^ or the bioprinting of sacrificial materials^20,65^ that create endothelial-lined channels hundreds of microns in diameter. By combining fabrication methods, one could create an engineered tissue that is initially perfused with larger endothelial-lined channels but includes densely patterned ECs branching from these channels. Over a few days, the patterned ECs could self-organize into microvessels that anastomose with the channels. This approach would combine the advantages of DPAC (microvascular resolution, complete control over structure) with the scalability and perfusability of top-down biofabrication methods.

An important limitation of this study is that the vessels were not continuously perfused during culture and were only perfusable after being pierced with a micropipette. Continuous perfusion might have further facilitated lumen formation^66^, prevented the buildup of luminal debris, and aided maturation and quiescence^67,68^. DPAC imposes several engineering constraints onto a potential microfluidic perfusion system, including the need to sandwich multiple layers of hydrogel and PDMS while remaining watertight. Incorporation of perfusion earlier in the self-organization process will require that these engineering challenges are overcome.

## Disclosures

- Z.J.G. is an equity holder in Scribe Biosciences and Provenance Bio.

## Author Contributions

K.A.C., V.S., and C.S. performed experiments. M.C.C. helped develop the methodology and contributed to the paper’s ideas. Z.J.G. and V.W. supervised experiments. K.A.C. and Z.J.G wrote the paper with feedback and approval from all authors.

## Acknowledgments

This research was supported in part by grants from the Department of Defense Breast Cancer Research Program (W81XWH-10-1-1023 and W81XWH-13-1-0221), NIH (U01CA199315, DP2 HD080351-01, 1R01CA190843-01, 1R21EB019181-01A, and 1R21CA182375-01A1), the NSF (MCB1330864), and the UCSF Center for Cellular Construction (DBI-1548297), an NSF Science and Technology Center. Z.J.G is a Chan-Zuckerberg BioHub Investigator. Sorting of fluorescent cells was done in the UCSF Laboratory for Cell Analysis, a core facility supported by a National Cancer Institute Cancer Center Support Grant (P30CA082103). Viruses were produced in the Viracore at UCSF.

---

**Figure S1:**
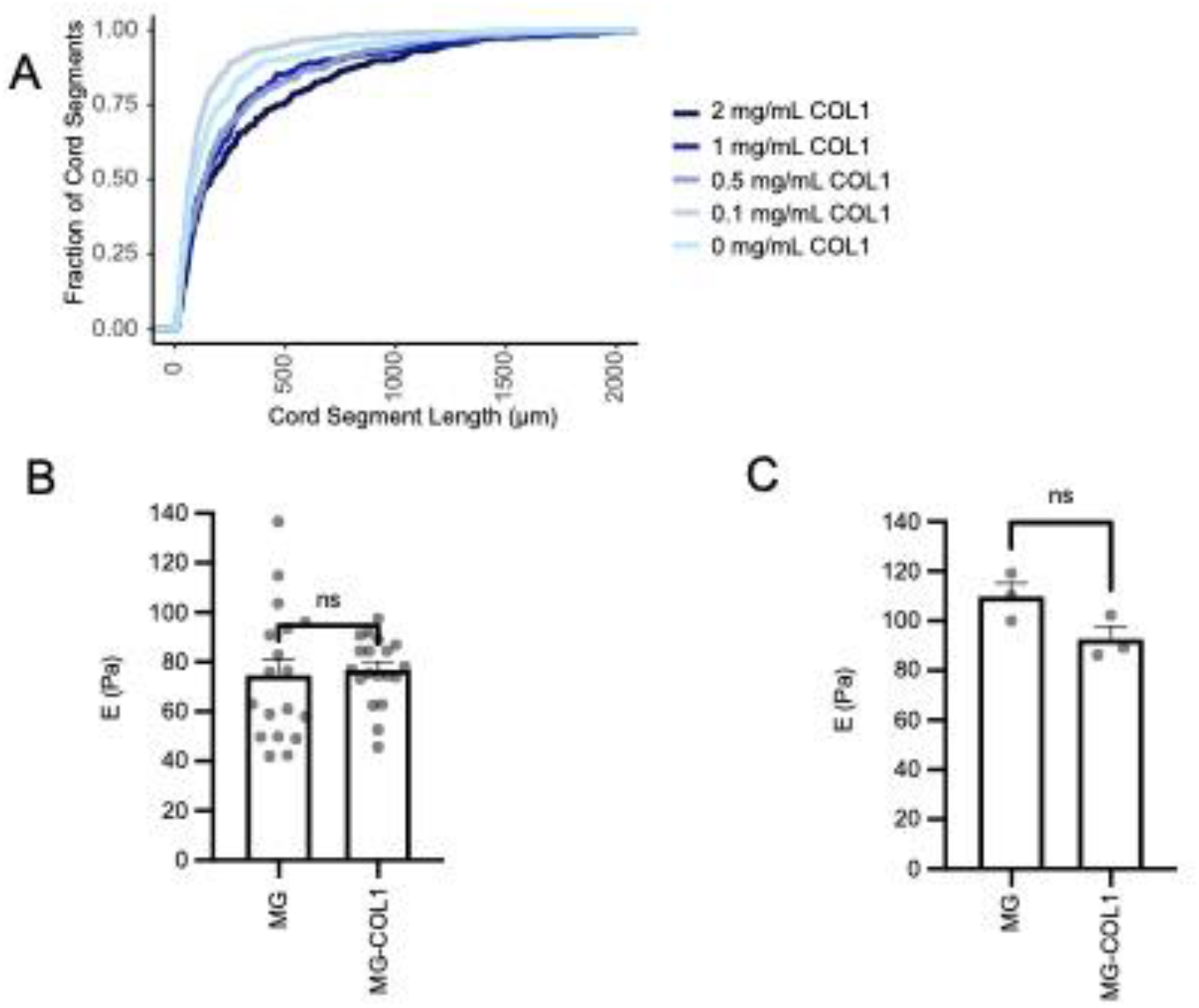
Collagen I contributes to the formation of longer continuous cord segments through a stiffness-independent mechanism. A) An empirical cumulative distribution function plot of the length of cord segments at 24h as a function of hydrogel composition. Hydrogels varied in the ratio of Matrigel to collagen I but maintained a total protein concentration of 8 mg/mL. B) Measurement of hydrogel stiffness (E) by rheology. T-test, not significant. C) Measurement of hydrogel stiffness (E) by atomic force microscopy. T-test, not significant.

**Figure S2:**
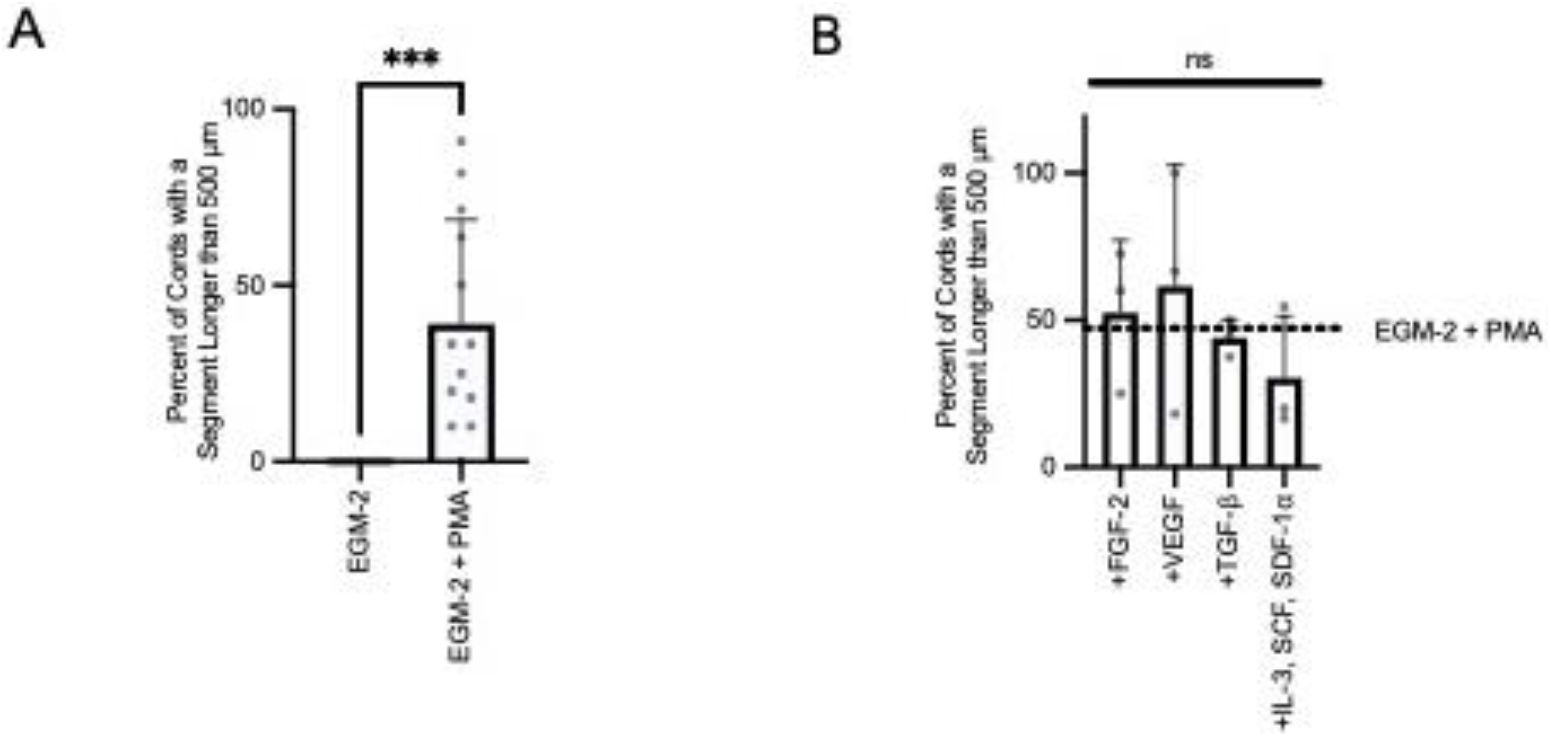
Pro-vasculogenic media components do not prevent scattering of HUVECs in patterned cords. A) The percent of cords per tissue that had at least one segment longer than 500 µm in length at 72 hours of culture in EGM-2 either containing or omitting 50 ng/mL PMA. Bars represent the mean ± SEM. Each data point represents one independent tissue. T-test, p<0.001. B) The percent of cords per tissue that had at least one segment longer than 500 µm in length at 72 hours of culture in EGM-2 + PMA with different media additives: 50 ng/mL VEGF, 50 ng/mL FGF-2, 50 ng/mL TGF-β, or a stem cell cytokine cocktail composed of 200 ng/mL each of SDF-1α, SCF, and IL-3. Bars represent the mean ± SEM. Each data point represents one independent tissue. One-way ANOVA with multiple comparisons to the control (EGM-2 + PMA), not significant.

**Figure S3:**
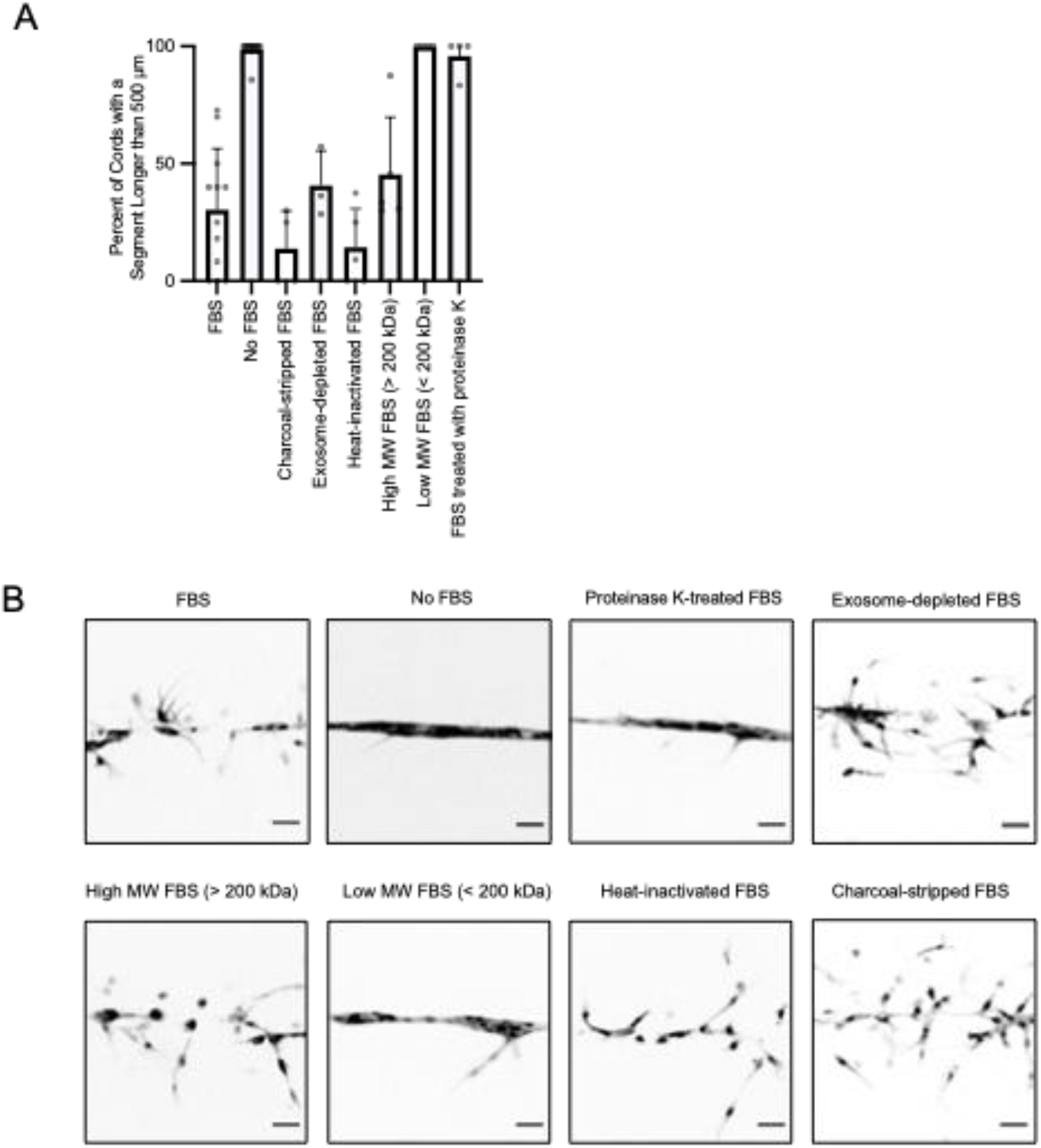
The component of FBS contributing to EC scattering contains a protein greater than 200 kDa. HUVEC cords were cultured in EGM-2 + PMA with the addition of one of several FBS variants, including charcoal-stripped FBS, exosome-depleted FBS, heat-inactivated FBS, FBS treated for 1 hour with 100 µg/mL proteinase K at 55°C before being neutralized with 100 µg/mL PMSF, FBS that has been filtered to exclude components greater than 200 kDa, and FBS that has been filtered to be enriched for components greater than 200 kDa. A) Analysis of the percent of cords per tissue that had at least one segment longer than 500 µm in length at 72 hours of culture as a function of media condition. Data represented as mean ± SEM with individual data points. All conditions were measured in at least three separate experiments. B) Representative images of cords cultured for 72h in different media formulations. Scale bar = 50 µm.

**Figure S4:**
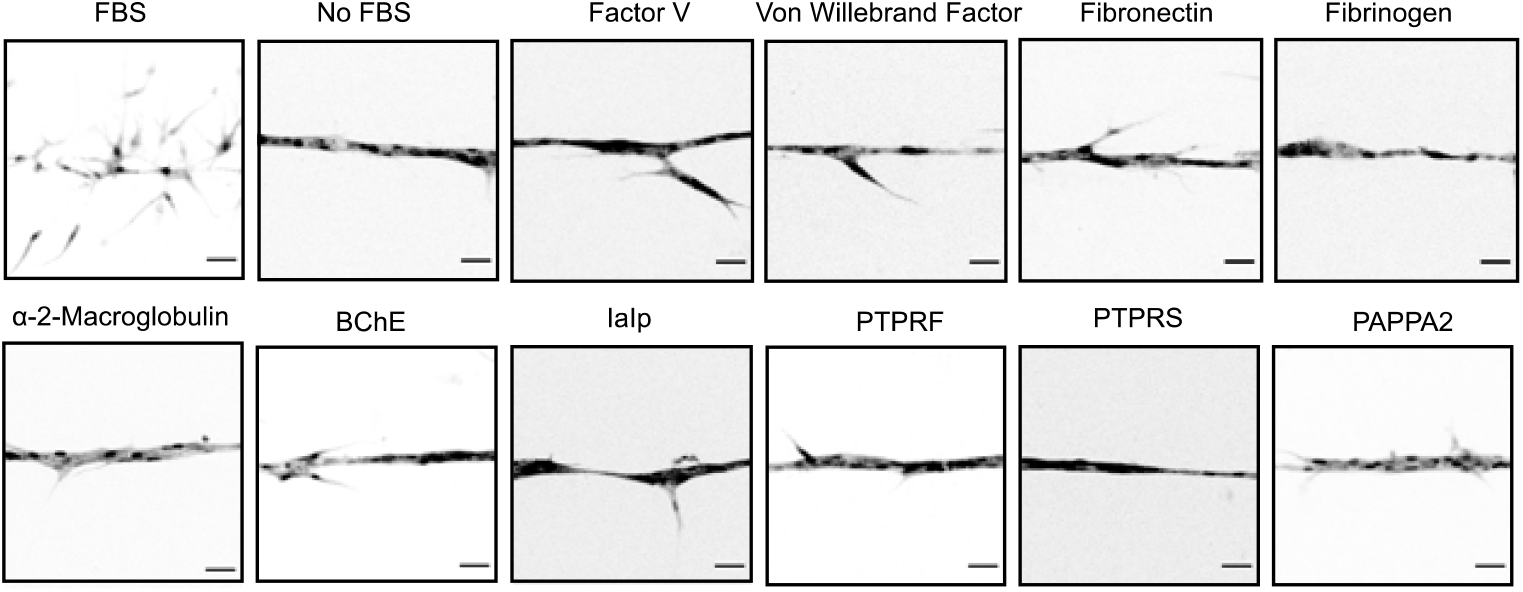
Phenotypes of candidate proteins in FBS larger than 200 kDa. Representative images of HUVEC cords cultured in serum-free media with the addition of recombinant proteins that fit the overall profile of being larger than 200 kDa and known to be present in serum.

**Figure S5:**
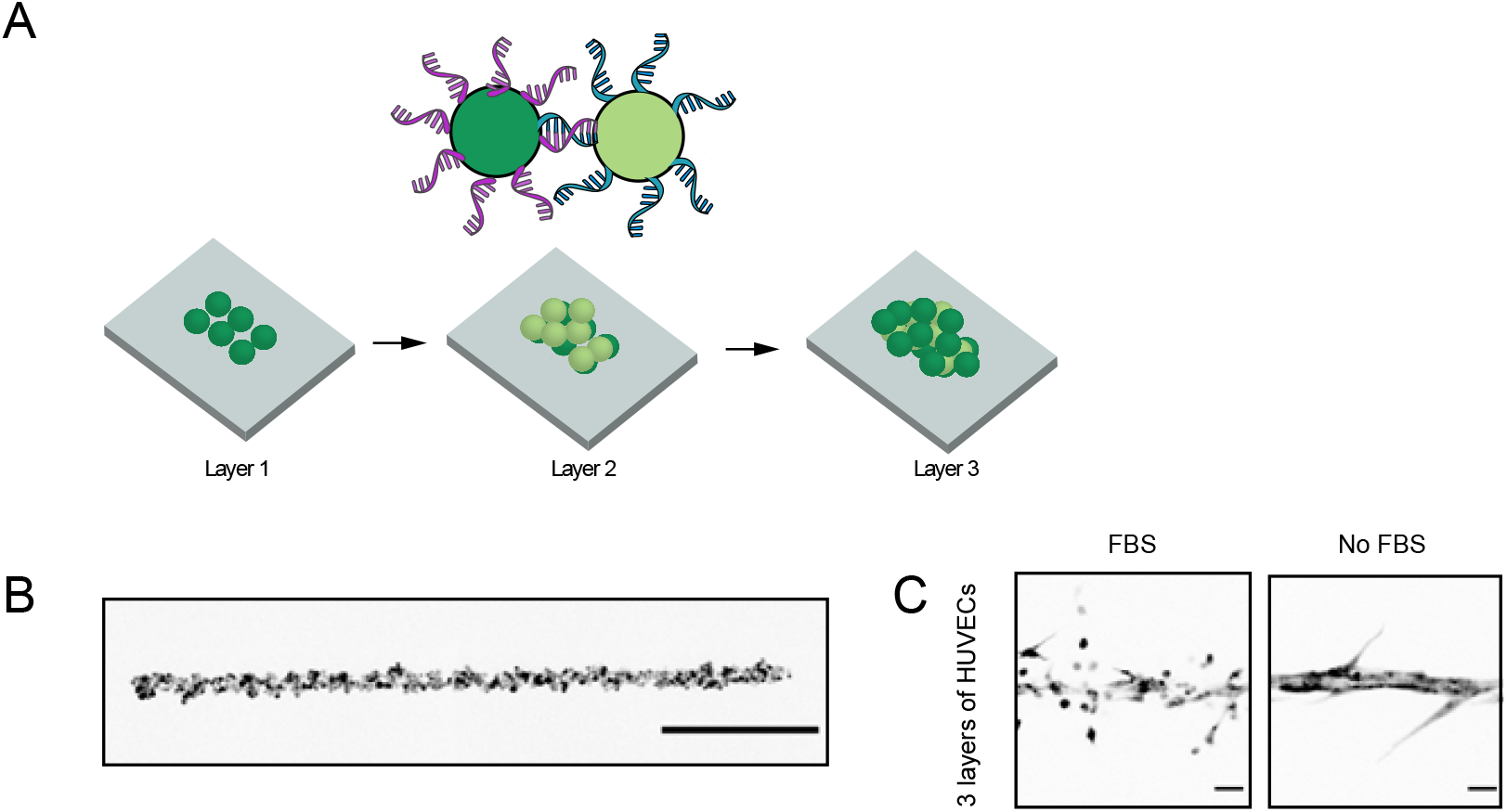
Multi-layered HUVEC cords respond to FBS with increased sprouting. A) Multi-layered cords can be created by first patterning one population of HUVECs as described in Figure S1, then adding a second population of HUVECs labeled with an oligo complementary to the first population of cells, then adding the first population of HUVECs again. Hybridization of the oligos dictates cell-cell adhesion, building up the tissue layer by layer. B) A three-layered HUVEC cord immediately after patterning. Scale bar = 500 µm. C) The response of three-layered HUVEC cords to FBS is like that of one-layered HUVEC cords. Scale bar = 50 µm.

**Figure S6:**
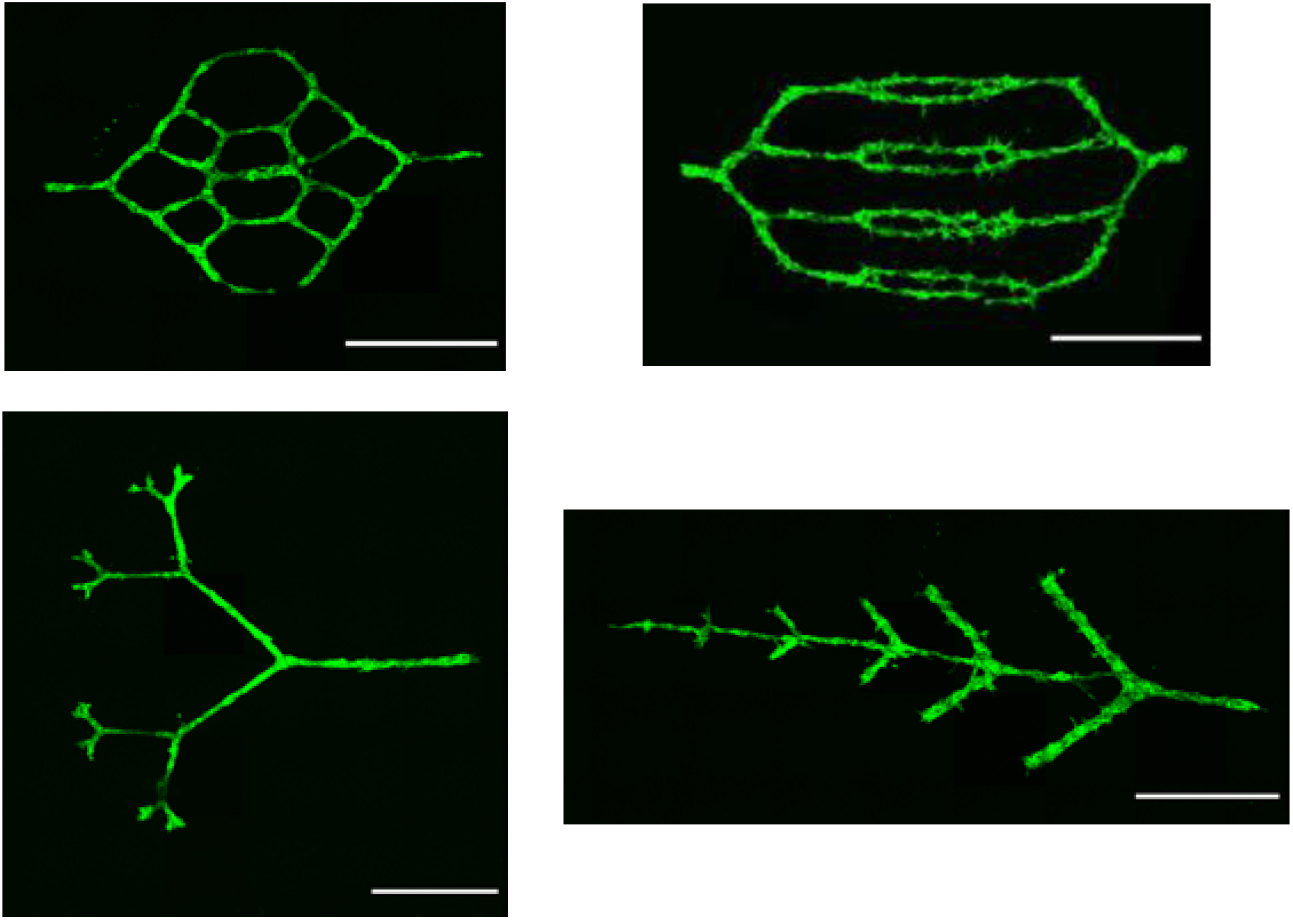
Branched microvascular structures created using DPAC. Hierarchical, branched microvascular structures millimeters in length can be designed and engineered using DPAC with complete control over the final structure. Pictured are patterned EGFP-HUVECs after 24 hours of culture. Scale bar: 1000 µm.

## Supplemental Video Figure Legends

**Video S1:** Cords composed of both EGFP-HUVECs (green) and mCherry-HUVECs (magenta) were cultured with EGM-2. Images were taken once per hour from 48h to 72h. Scale bar: 50 µm.

**Video S2:** Cords composed of both EGFP-HUVECs (green) and mCherry-HUVECs (magenta) were cultured with EGM-2 without FBS. Images were taken once per hour from 48h to 72h. Scale bar: 50 µm.

**Video S3:** Pairs of EGFP-HUVECs were patterned 25 µm apart (center-to-center distance) and cultured in FBS-containing EGM-2 + PMA for 70 hours and imaged once per hour. Scale bar: 100 µm.

**Video S4:** Pairs of EGFP-HUVECs were patterned 25 µm apart (center-to-center distance) and cultured in FBS-free EGM-2 + PMA for 70 hours and imaged once per hour. Scale bar: 100 µm.

**Video S5:** Perfusion of a 3-layered HUVEC microvessel cultured without FBS for 7 days. The microvessel was cut open on one end using a scalpel to create an exit for the perfused fluid. Injection of PBS with an aluminosilicate glass micropipette pushed debris out of the microvessel, revealing an intact lumen. Scale bar: 100 µm.

**Video S6:** Perfusion of a 3-layered mCherry-HUVEC microvessel cultured without FBS for 7 days with a solution of 2 µm diameter blue, fluorescent beads. Scale bar: 100 µm.

**Video S7:** Perfusion of a 3-layered branched microvascular network cultured without FBS for 7 days. Injection of PBS into the central branch clears the debris through all branches, indicating a continuous open lumen throughout the structure.

## Notes

### Competing Interest Statement

Zev Gartner is an equity holder in Scribe Biosciences and Provenance Bio.

