## Supplemental Methods for "Microenvironmental engineering improves the self-organization of patterned microvascular networks"

### Creation of PDMS Flow Cells:

Rectangular PDMS flow cells (dimensions either 10 mm x 15 mm x 200 µm or 4 mm x 19 mm x 200 µm, depending on experiment) are used to concentrate the cells over the patterned regions of the slide and serve as a mold for the hydrogel. SU-8 master wafers for the flow cells were created by spin coating SU-8 2075 (Microchem) onto a silicon wafer (500 rpm for 10s, followed by 1000 rpm for 30s) to create features approximately 200 µm in height.[31] The wafer was baked at 95 °C for 45 minutes to remove excess solvent. The wafer was then exposed to UV light (365 nm) traveling through a photomask for a radiant energy density of 350 $mJ/cm^2$. The wafer was then baked at 95°C for 15 minutes. The wafer was developed by immersion in SU-8 Developer Solution (PGMEA) for 15 minutes. After development was complete, the wafer was dried and baked for 5 additional minutes.

Polydimethylsiloxane (PDMS) was prepared by mixing polydimethylsiloxane elastomer (Sylgard 184, Corning) and crosslinker in a 10:1 ratio by mass. The PDMS solution was de-gassed in a vacuum desiccator for 15 minutes to remove bubbles before it was poured onto the master wafer. The PDMS was cured by baking in a 60 °C oven for 3 hours.

Shortly before starting a DPAC experiment, the required number of PDMS flow cells were cut out from the master wafer. Plasma oxidization with 20 cc/min room air for 5 minutes rendered the surfaces of the flow cells hydrophilic. The flow cells were cut to have 1-2 mm of PDMS remaining on each side and then the top and bottom of the flow cell were cut open to create an inlet and outlet. Using the photomask as a reference, PDMS flow cells were placed on top of the slide in the location of each patterned region.

**Rheology:**

Shear rheology data was collected using an AR 2000ex rheometer (TA Instruments). 300 µL gels were deposited onto the base of the rheometer, which was heated to 37 °C. Then, a 25 mm flat plate probe was lowered, and the sample was compressed into a barrel. Approximately 250 µL of mineral oil (Sunmark™) was applied around the perimeter of the sample to prevent evaporation. Time sweep data was collected with 1% strain at a frequency of 1 rad/s until the loss and storage moduli reached steady-state values, approximately 30 minutes to an hour. The elastic modulus, E, was calculated by the formula $E = 2(1 + v) G^*$, where Poisson's ratio $v$ was assumed to be 0.5 and where $G^*$ is the complex modulus found using the measured storage and loss moduli: $G^* = (G'^2 + G''^2)^{1/2}$.[36]

**Atomic Force Microscopy:**

For AFM measurements, 300 µL gels were deposited directly onto glass slides. 15 mm round coverslips coated in a thin layer of hydrophobic Rain-X (R) solution were immediately placed on top to flatten the gels. Coverslips were removed after polymerization at 37 °C for one hour. Then, the surrounding areas were marked with a hydrophobic barrier and gels were submerged in approximately 500 µL of PBS. Measurements were obtained with a MFP3D-BIO inverted optical AFM mounted on a Nikon TE2000-U inverted fluorescent microscope (Asylum Research). Gels were indented with silicon nitride cantilevers with borosilicate glass spherical tips 5 um in diameter (Novascan Tech). Cantilevers had a nominal spring constant of 0.06 N/m and were calibrated using the thermal oscillation of the tip. Gels were indented at 2 µm/s loading rate with a 1.8 nN force trigger. Gel stiffness was computed using the Hertz model with a sample Poisson Ratio of 0.5.

**Antibodies used:**

Primary antibodies used: VE-cadherin (ab33168, Abcam), ZO-1 (ZO-1 Monoclonal Antibody, Invitrogen), collagen IV (ab6586, Abcam), claudin-5 (Claudin 5 Monoclonal Antibody (4C3C2), Invitrogen), laminin α5 (ab14509, Abcam).

Secondary antibodies used: Goat anti-Mouse IgG (H+L) Cross-Adsorbed Secondary Antibody, Alexa Fluor 488 (Invitrogen), Goat anti-Mouse IgG (H+L) Cross-Adsorbed Secondary Antibody, Alexa Fluor 647 (Invitrogen), Goat anti-Rabbit IgG (H+L) Cross-Adsorbed Secondary Antibody, Alexa Fluor 405 (Invitrogen), Goat anti-Rabbit IgG (H+L) Cross-Adsorbed Secondary Antibody, Alexa Fluor 488 (Invitrogen), Goat anti-Rabbit IgG (H+L) Cross-Adsorbed Secondary Antibody, Alexa Fluor 568 (Invitrogen), Goat anti-Rabbit IgG (H+L) Cross-Adsorbed Secondary Antibody, Alexa Fluor 647 (Invitrogen).

DAPI was used to visualize nuclei.

**FBS candidate protein experiments:**

Candidates for the active component of FBS were proteins larger than 200 kilodaltons in mass, known to be present in serum, and known in the literature to be associated with endothelial cells or angiogenesis. Recombinant versions of these proteins were added into serum-free EGM-2 at the concentrations listed below.

| Candidate Protein | Molecular Weight | Concentration in Media | Supplier |
|---|---|---|---|
| Alpha-2-macroglobulin | 800 | 44 µg/mL | Sigma-Aldrich |
| Apolipoprotein B | 240 | 20 µg/mL | Sigma-Aldrich |
| Butyrylcholinesterase | 260 | 100 ng/mL | R&D Systems |
| Factor V (bovine) | 330 | 200 ng/mL | Enzyme Research Laboratories |
| Fibrinogen | 340 | 40 µg/mL | Sigma-Aldrich |
| Fibronectin | 220 | 5 µg/mL | Sigma-Aldrich |

| | | | |
|---|---|---|---|
| Inter Alpha Inhibitor Protein | 225 | 10 µg/mL | Athens Research and Technology |
| PAPP-A2 | 220 | 0.5 ng/mL | R&D Systems |
| PTPRF | 213 | 200 ng/mL | R&D Systems |
| PTPRS | 217 | 200 ng/mL | Sigma-Aldrich |
| VWF | 500 | 10 µg/mL | EMD Millipore |

**Oligonucleotide Sequences for Cell Labeling:**

Cells were either labeled with lipid-modified oligos (LMOs) or cholesterol-modified oligos (CMOs) for the purposes of cell patterning. We have previously determined that the CMOs and LMOs perform equally well for cell patterning[34]. We transitioned from the LMOs to the CMOs because the CMOs are inexpensive and do not require in-house synthesis and purification.

For lipid-modified oligo labeling:

Surface Oligo: 5' (Amine C6) ACTGACTGACTGACTGACTG 3'

Cell Oligo: 5' (Lignoceric Acid) - GTA ACG ATC CAG CTG TCA CT TTTTTTTTTT TTTTTTTTTT TTTTTTTTTT TTTTTTTTTT TTTTTTTTTT TTTTTTTTTT CAGT CAGT CAGT CAGT CAGT

Co-Anchor Oligo (for stabilization)[38]: 5' AGT GAC AGC TGG ATC GTT A – (Palmitic Acid) 3'

For cholesterol-modified oligo labeling:

Surface Oligo: 5' (Amine C6) ACTGACTGACTGACTGACTG 3'

Universal Anchor: 5'-TGAATCTCTGGGTGCCAAGGGTAAGCATCCAGCTGTCACT - {Chol}-3'

Universal Co-Anchor: 5'-{Chol} AGTGACAGCTGGATGCTTAC -3'

Adapter Strand A prime: CCTTGGCACCCAGAGATTCA TTTTTTTTTTTTTTTTTTTT CAGTCAGTCAGTCAGTCAGT
